# Loss of BICD2 in muscle drives motor neuron loss in a developmental form of spinal muscular atrophy

**DOI:** 10.1101/854711

**Authors:** AM Rossor, JN Sleigh, M Groves, F Muntoni, MM Reilly, CC Hoogenraad, G Schiavo

**Author notes:** Corresponding authors: Alexander M Rossor:, Giampietro Schiavo.

## Abstract

BICD2 is a key component of the dynein/dynactin motor complex. Autosomal dominant mutations in *BICD2* cause Spinal Muscular Atrophy Lower Extremity Predominant 2 (SMALED2), a developmental disease of motor neurons. In this study we sought to examine the motor neuron phenotype of conditional *Bicd2*^−/−^ mice. *Bicd2*^−/−^ mice show a significant reduction in the number of motor axons of the L4 ventral root compared to wild type mice. Muscle-specific knockout of *Bicd2*, but not motor neuron-specific *Bicd2* loss, results in a reduction in L4 ventral axons comparable to global *Bicd2*^−/−^ mice. Rab6, a small GTPase required for the sorting of secretory vesicles from the TGN to the plasma membrane is a major binding partner of BICD2. We therefore examined the secretory pathway in SMALED2 patient fibroblasts and demonstrated impaired flow of constitutive secretory cargoes. Together, these data indicate that BICD2 loss from muscles is a major driver of non-cell autonomous pathology with important implications for future therapeutic approaches to SMALED2.

**Summary:** Missense mutations in the cargo adaptor protein BICD2 cause SMALED2, a developmental disease of motor neurons. In this study, the authors show that *BICD2* mutations cause motor neuron loss by a non-cell autonomous mechanism determining a disabling impairment of muscle function.

## Introduction

Spinal Muscular Atrophy Lower Extremity Predominant (SMALED) is a disease of lower motor neurons, principally affecting the lower limbs. Affected individuals often present at birth with contractures of the lower limbs. Autosomal dominant missense mutations in *DYNC1H1* and *BICD2* are the only two known genetic causes of SMALED and determine two indistinguishable forms of this disease (Oates et al., 2013; Neveling et al., 2013; Scoto et al., 2015; Harms et al., 2012; Peeters et al., 2013).

*DYNC1H1* encodes the cytoplasmic dynein heavy chain, a key component and force-generating subunit of the dynein/dynactin retrograde transport complex. This complex is responsible for the transport of intracellular cargoes towards the minus-end of microtubules located in the cell soma (Schiavo et al., 2013). In contrast, anterograde transport towards the positive end of microtubules is driven by the kinesin family of motor proteins (Klinman and Holzbaur, 2018).

Motor neurons have extremely long axons, which make them preferentially susceptible to deficits in axonal transport (Sleigh et al., 2019). It has long been thought that mutations in *BICD2* and *DYNC1H1* impair axonal transport leading to motor neuron degeneration. This is supported by work in *Drosophila* demonstrating a reduction of the *in vitro* run length in flies expressing disease-causing mutant DYNC1H1 compared to wild type (Hoang et al., 2017). This conclusion has been brought into question by the contrasting findings that several disease-causing mutations in *BICD2* increase its binding affinity to the dynein/dynactin complex resulting in an increase in run length (Huynh and Vale, 2017).

BICD2 is a cargo adaptor protein, comprising a N-terminal region that mediates binding to the dimerization domain of DYNC1H1 and a C-terminal cargo binding domain (Hoogenraad et al., 2001). Missense mutations throughout *BICD2* have been shown to cause SMALED2 (Storbeck et al., 2017), yet the disease mechanism is unclear. Mutations in the N-terminal domain increase the affinity to the dynein complex (Huynh and Vale, 2017), whereas the p.Glu774Gly mutation in the C-terminal domain has no effect on DYNC1H1 binding, but disrupts the interaction with its cargo Rab6, causing a loss of function phenotype (Peeters et al., 2013). As BICD2 forms dimers, such mutations may also impair the cargo binding ability of wild type/mutant BICD2 heterodimers, leading to near complete loss of function as opposed to haploinsufficiency. A loss of function effect in SMALED2 was thought to be unlikely as the *Bicd2*^−/−^ mouse was originally reported to lack a motor phenotype (Jaarsma et al., 2014). However, recent studies of SMALED2 patients with severe mutations in *BICD2* have revealed additional cortical and cerebellar phenotypes similar to the *Bicd2*^−/−^ mouse. These findings suggest that pathological *BICD2* mutations may indeed induce a loss of function (Ravenscroft et al., 2016). In this study, we sought to re-examine the motor neuron phenotype in the *Bicd2*^−/−^ mouse to conclusively assess the molecular basis of the pathomechanism of SMALED2.

## Results and Discussion

### *Bicd2*^−/−^ mice display a significant motor neuron loss

Due to the similarities between severe SMALED2 patients and *Bicd2*^−/−^ mice (Ravenscroft et al., 2016; Jaarsma et al., 2014) and the observation that missense mutations in the cargo binding domain of BICD2 impair Rab6 binding with no effect on dynein binding (Peeters et al., 2013), we predicted a loss of function pathomechanism in SMALED2. Following this hypothesis, loss of BICD2 function should result in motor neuron loss. We therefore re-examined the motor neuron phenotype of *Bicd2*^−/−^ mice by examining the L4 dorsal and ventral nerve roots in the spinal cord of wild type and knockout mice at postnatal day 21 (p21). We were unable to examine these mice at later time points as knockout mice die by four weeks of age from obstructive hydrocephalus (Jaarsma et al., 2014). The L4 nerve root was chosen as it is the largest lumbar nerve root. At p21, there is a significant reduction in the total number of motor axons in *Bicd2*^−/−^ mice compared to wild type and this is restricted to a subpopulation of motor axons with a diameter of 2.5-4 μm (**Figure 1**; *Bicd2*^+/+^ mean = 926±22.7 (SEM, n=7), *Bicd2*^−/−^ = 806±25.0 (n=7), unpaired *t*-test, p=0.0041). In contrast, there was no reduction in the number of L4 dorsal root sensory axons between wild type (1,778±35.8; n=6) and *Bicd2*^−/−^ (1,792±60.4; n=6) mice (unpaired t-test, p=0.85). The lack of a sensory phenotype was corroborated by further analyses of the percentage of medium-to-large, NF200^+^ neurons, and small peripherin^+^ neurons, in the L4 dorsal root ganglia (DRG), which did not reveal any difference between wild type and *Bicd2*^−/−^ mice (**Supplementary Figure 1**).

**Figure 1.**
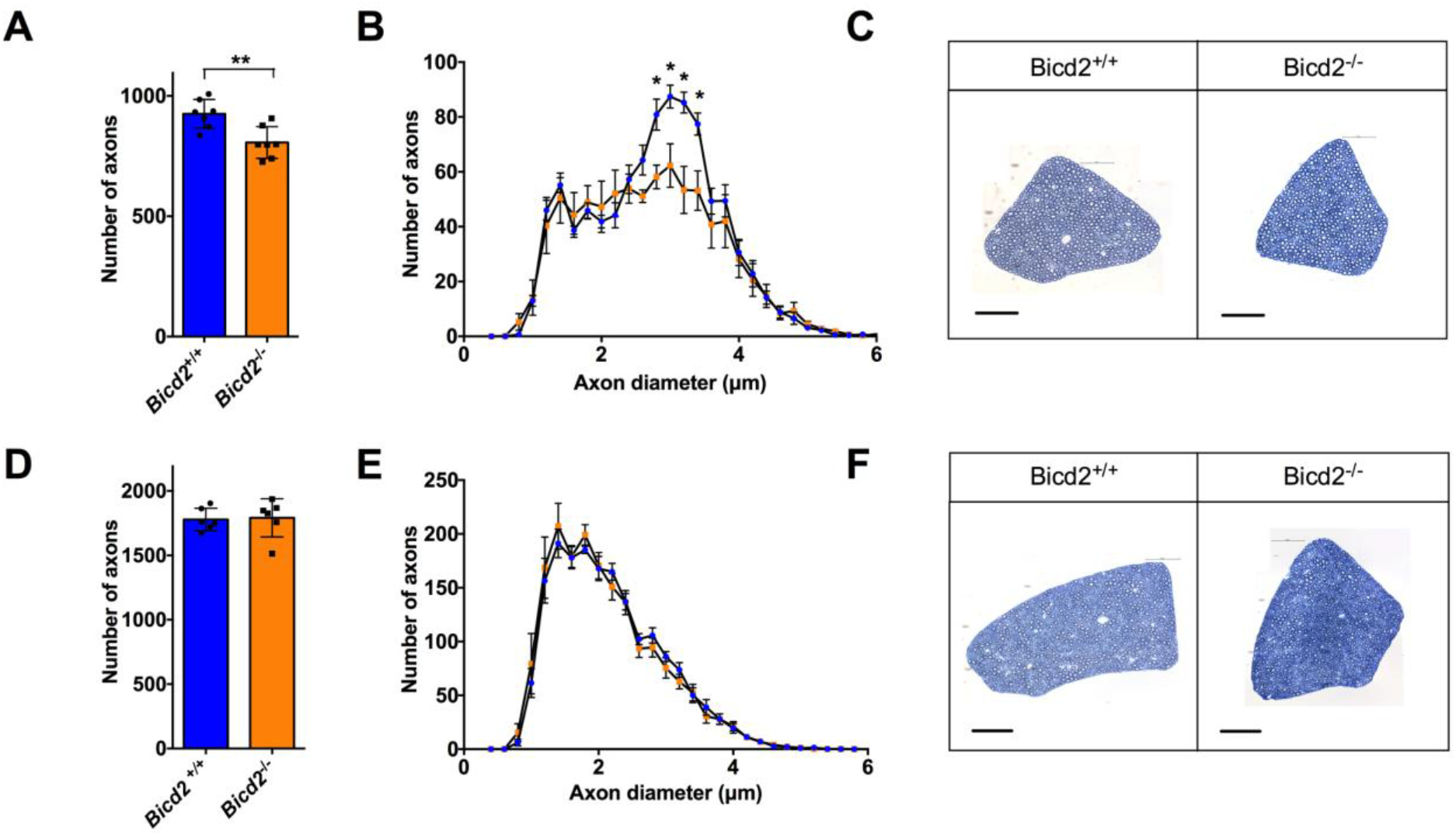
Loss of motor axons in *Bicd2*^−/−^ mice at 21 days of age. (**A**) shows the number of motor axons in the L4 ventral root of *Bicd2*^+/+^ (wild type) and *Bicd2*^−/−^ (knockout) mice (**unpaired 2-sided *t*-test, *p*=0.0041). (**B**) shows a histogram of the total number of L4 motor axons classified using 0.2 μm bins (blue=*Bicd2^−/−^*, orange=*Bicd2^−/−^*), \**p*<0.001; multiple *t*-tests corrected for multiple comparisons using the Holm-Sidak method. (**C**) Representative images from 0.8 μm cross-sections of 1% toluene blue stained L4 ventral roots of *Bicd2*^+/+^ and *Bicd2*^−/−^ mice. (**D**) shows the number of sensory axons in the L4 dorsal root of *Bicd2*^+/+^ and *Bicd2*^−/−^ mice. (**E**) shows a histogram of the total number of L4 sensory axons classified as in **B**(blue=*Bicd2*^−/−^, orange=*Bicd2*^−/−^). (**F**) Representative images from 0.8 μm cross sections of 1% toluene blue stained L4 dorsal roots of *Bicd2*^+/+^ and *Bicd2*^−/−^ mice. Error bars = standard error of the mean (SEM). Scale bars = 50 μm.

### Loss of motor neurons in *Bicd2*^−/−^ mice is caused by a muscle non-cell autonomous process

The cerebellar hypoplasia phenotype observed in *Bicd2*^−/−^ mice is due to a non-cell autonomous process arising in Bergmann glia (Jaarsma et al., 2014). Therefore, we hypothesised that the motor neuron loss we observed in the same knockout mice may also be due to a non-cell autonomous mechanism, but in this instance, arising from muscle tissue, which has previously been shown to have important trophic support functions for motor neurons (Kablar and Belliveau, 2005). According to the neurotrophin hypothesis, during early development, an excess of motor neurons reach their target muscle and compete for muscle-secreted survival factors (e.g. neurotrophins) (Kanning et al., 2010). A stochastic process of programmed cell death then follows, as those motor neurons that receive insufficient neurotrophins undergo apoptosis. To test this hypothesis, we generated mice in which *Bicd2* is selectively knocked out only in skeletal muscle. Homozygous *Bicd2* mice, in which the endogenous *Bicd2* allele is flanked by intronic loxP sequences (Jaarsma et al., 2014), were crossed with mice expressing cre-recombinase driven by the endogenous *Myod* promoter to generate mice lacking both *Bicd2* alleles in muscle tissue alone (Kanisicak et al., 2009; Jaarsma et al., 2014). These mice had a normal life span and no obvious gait abnormality. Analyses of these mice at p21 shows a significant reduction in motor axons in the L4 ventral nerve root compared to wild type mice (**Figure 2**; *Bicd2*^+/+^ = 928±24 (n=7), *Myod-Cre* = 817±39 (n=7), one-way ANOVA *p*=0.019, Dunnett’s *t*-test, *p*=0.034). Furthermore, the subpopulation of motor axons affected is the same as the *Bicd2*^−/−^ mouse (2.5-4.4 μm diameter). A small reduction in the number of L4 motor axons between wild type (928±24) and homozygous *Bicd2*^loxP/loxP^ (888±23, n=7; Dunnett’s *t*-test, *p*=0.705) mice was detected, but it did not reach statistical significance (**Figure 2A**). To confirm that the loss of motor axons in *Bicd2*^−/−^ mice was solely due to loss of Bicd2 in muscle, we generated mice lacking Bicd2 in motor neurons. Homozygous *Bicd2*^*loxP*/*loxP*^ mice were crossed with mice expressing cre-recombinase driven by the endogenous *ChAT* promoter to generate mice lacking both *Bicd2* alleles selectively in motor neurons. Analysis of the number of L4 motor axons in these genetically modified mice at p21 revealed no difference compared to Bicd2^−/−^ mice (**Figure 2**; *Bicd2*^+/+^ = 928±24; n=7; *ChAT-Cre* = 924±23, n=3; one-way ANOVA *p*=0.019, Dunnett’s *t*-test, *p*=0.939)

**Figure 2.**
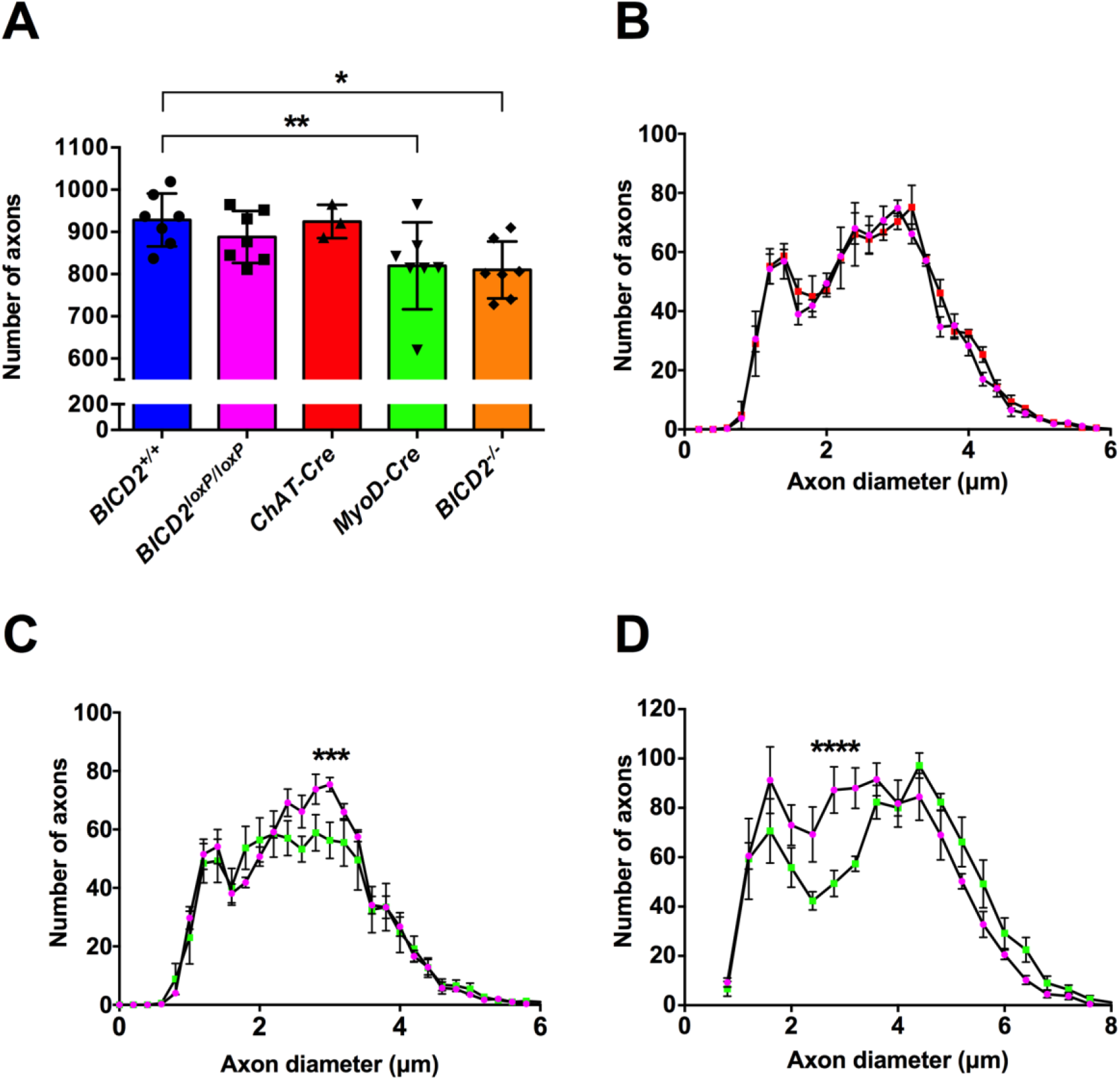
Motor axon loss in muscle specific *Bicd2* knockout mice. (**A**) shows the total number of axons at 21 days of age in *Bicd2*^+/+^ mice (blue), *Bicd2*^*loxP*/*loxP*^ mice in which both *Bicd2* alleles are flanked by loxP sites (pink), mice with motor neuron specific *Bicd2* knockout (*ChAT-Cre*, red), mice with muscle-specific *Bicd2* knockout (*Myod-Cre*, green), and *Bicd2*^−/−^ mice (orange). A significant reduction in the total number of L4 ventral axons was found in *Myod-Cre* and *Bicd2*^−/−^ mice compared to wild type (one-way ANOVA *p*=0.0187, Dunnett’s 2-sided *t*-test with *Bicd2*^+/+^ as control, \*\**p*=0.019, \**p*=0.034). (**B**) shows a histogram of the total number of L4 ventral axons at 21 days of age classified using 0.2 μm bins (pink=*Bicd2*^*loxP*/*loxP*^, red=*ChAT-Cre*). (**C**) shows a histogram of the total number of L4 motor axons at 21 days of age classified using 0.2 μm bins (pink=*Bicd2*^*loxP*/*loxP*^, green=*Myod-Cre*); multiple *t*-tests corrected for multiple comparisons using the Holm-Sidak method, \*\*\**p*=0.0009. (**D**) shows a histogram of the total number of L4 motor axons at 42 days of age classified using 0.4 μm bins (pink= *Bicd2*^*loxP*/*loxP*^ (n=4), green=*Myod-Cre* (n=6); multiple t-tests corrected for multiple comparisons using the Holm-Sidak method,\*\*\*\**p*=0.0002). Error bars = SEM.

A similar analysis performed on the ‘legs at odd angles mouse’ (loa) strain, which bear a missense mutation in *Dync1h1* (a binding partner of *Bicd2*), showed at a later time point of six weeks a reduction in small diameter motor axons, which were assumed to be gamma motor neurons (Hafezparast et al., 2003; Ilieva et al., 2008). We therefore repeated this analysis in *Myod-Cre* mice at six weeks of age. A comparison of the axon diameter cumulative distribution curves shows a loss of motor axons in mice with muscle-specific knockout of *Bicd2*, revealing a decrease in a subset of motor axons of smaller diameter in the putative range of gamma motor neurons (**Figure 2C**).

### Active denervation is absent in *Bicd2*^−/−^ mice

Next, we sought to determine whether the loss of motor neurons was due to active degeneration of motor axons. Morphological analysis of semi-thin ventral root sections revealed no acute axon degeneration profiles (**Figure 1**). We therefore analysed the neuromuscular junctions (NMJs) from two different muscles (lumbricals and flexor digitorum brevis (FDB) of the hind paw). Wholemount preparations of these two thin muscles were chosen due to reliable quantification of NMJ degeneration without the need for sectioning (Sleigh et al., 2014). Interestingly, all muscles showed a full NMJ innervation pattern, suggesting that at the time of examination, there was no significant denervation (**Figure 3A** **and** **C**). Furthermore, there was no evidence to indicate that the post-natal developmental process of synapse elimination at the NMJ was affected in *Bicd2*^−/−^ mice (**Figure 3B**). In contrast, the NMJ area of the hindfoot lumbrical and FDB muscles was significantly smaller in *Bicd2*^−/−^ compare to wild type mice (**Figure 3D**). The significance of this is not clear but may be a consequence of the significant reduction in overall size of *Bicd2*^−/−^ mice and their muscle mass compared to wild type animals (**Supplementary Figure 2A and C**).

**Figure 3.**
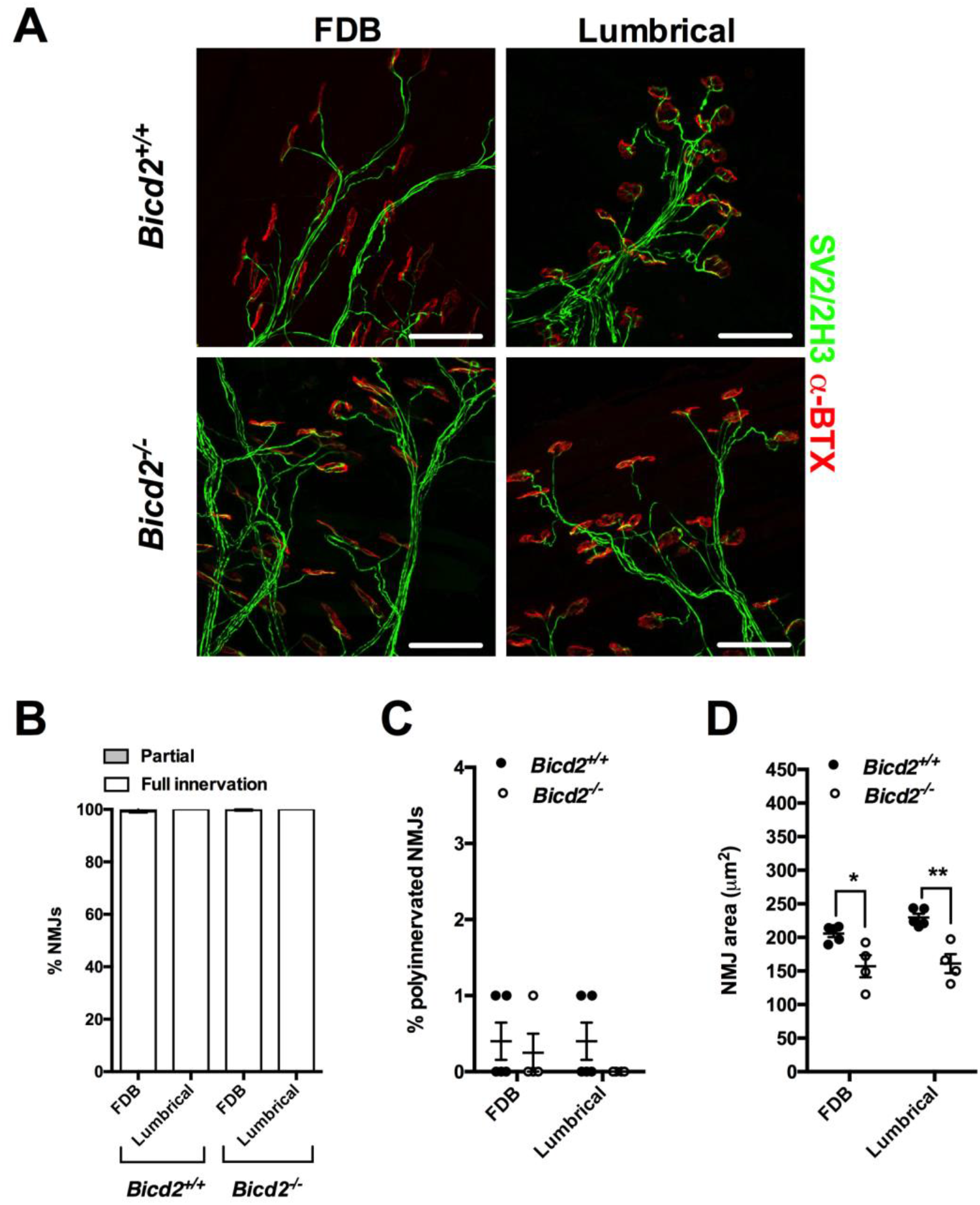
Normal NMJ analysis of *Bicd2*^−/−^ mice at 21 days of age. (**A**) shows representative images of the NMJs of the FDB (flexor digitorum brevis) and feet lumbrical muscles stained with anti-SV2/2H3 antibodies (green) to visualise motor neurons and fluorescent alpha-bungarotoxin (red) to identify post-synaptic acetylcholine receptors on the muscle fibre surface. Scale bars = 50 μm. (**B**) shows the percentage of fully and partially innervated NMJs in *Bicd2*^+/+^ (n=4) and *Bicd2*^−/−^ (n=4) mice. (**C**) shows the percentage of poly-innervated (measure of immaturity) NMJs between *Bicd2*^+/+^ (n=4) and *Bicd2*^−/−^ (n=4) mice. (**D**) shows the area of the NMJ in the FDB and lumbrical muscles in *Bicd2*^+/+^ (mean 205 and 230 μm^2^, respectively; n=4) and *Bicd2*^−/−^ (mean 157 and 161 μm^2^, respectively; n=4) mice, (multiple *t*-tests corrected for multiple comparisons using the Holm-Sidak method, \**p*=0.05, \*\**p*=0.004). Error bars = SEM.

### Muscles in *Bicd2*^−/−^ mice display loss of muscle spindles and gamma motor neurons, but no gross morphological changes

Patients with SMALED2 show muscle biopsy abnormalities indicative of a primary muscle pathology (Unger et al., 2016). We therefore examined the gastrocnemius muscle of wild type and *Bicd2*^−/−^ mice, but did not identify any consistent differences on H&E-stained muscle or on electron microscopy (**Supplementary Figure 3**).

Motor neuron subtypes may be subdivided according to the fibre type they innervate (Kanning et al., 2010). For example, ‘fast’ alpha motor neurons innervate types IIa, IIb and IIx muscle fibres, whereas slow motor neurons innervate type 1 muscle fibres. We therefore examined the proportion of muscle fibres of the L4 innervated gastrocnemius muscle in wild type and *Bicd2*^−/−^ mice, but we found no difference (**Supplementary Figure 2**), indicating that *Bicd2* ablation does not induce a significant muscle fibre switch.

As shown in **Figure 2**, at both three and six weeks of age, the population of motor axons lost in the L4 ventral roots of mice in which *Bicd2* has been selectively knocked out in muscle falls within the 2.5-4 μm range also seen in Bicd2^−/−^ mice. The lack of denervation of NMJs or switch in muscle fibre types, suggests that alpha motor neurons are not affected. At six weeks of age, based on axon diameter, the loss of axons falls within the (**Figure 2**) presumed gamma motor neuron population. This is the same population of motor neurons that is lost in mice with missense mutations in *Dync1h1* (Ilieva et al., 2008), which model SMALED in humans. Gamma motor neurons represent approximately 30% of motor neurons innervating muscle, but, unlike large calibre alpha motor neurons, they do not form NMJs with skeletal muscle fibres and instead contact muscle spindles. At three weeks, the distinction of motor neuron subtype based on axon diameter is unreliable (Kanning et al., 2010). To confirm whether the loss of L4 ventral axons at 3 weeks also correlated with a reduction in gamma motor neurons, we examined the total number of muscle spindles in the soleus muscle. The soleus muscle was chosen as it is a small muscle in which the total number of muscle spindles can be accurately quantified on serial transverse sections as previously described (Sleigh et al., 2017). By staining for the SV2 and 2H3 antigens, muscle spindles can be identified, although gamma motor neuron efferents and 1a sensory afferents are indistinguishable. A comparison of the total number of muscle spindles revealed a 20% reduction in *Bicd2*^−/−^ mice (9.4±0.68) compared to wild type (12.5±0.29, unpaired *t*-test *p*=0.0065) (**Figure 4**). This correlates well with the reduction in smaller diameter motor axons at six weeks in *Myod-Cre* mice (**Figure 2C**) and confirms that the motor axons lost at three weeks are also gamma motor neurons. The innervation pattern of the remaining muscle spindles was unperturbed, suggesting a developmental, as opposed to a degenerative process. Of note, there was no reduction in large diameter (presumed sensory 1a afferents) in the L4 dorsal nerve root of *Bicd2*^−/−^ (**Figure 1D-E**) and *Myod-Cre* mice (data not shown). It is possible that SV2 staining is specific to the gamma motor neuron endplate and does not stain for sensory 1a afferents that do not form synapses at the muscle spindle.

**Figure 4.**
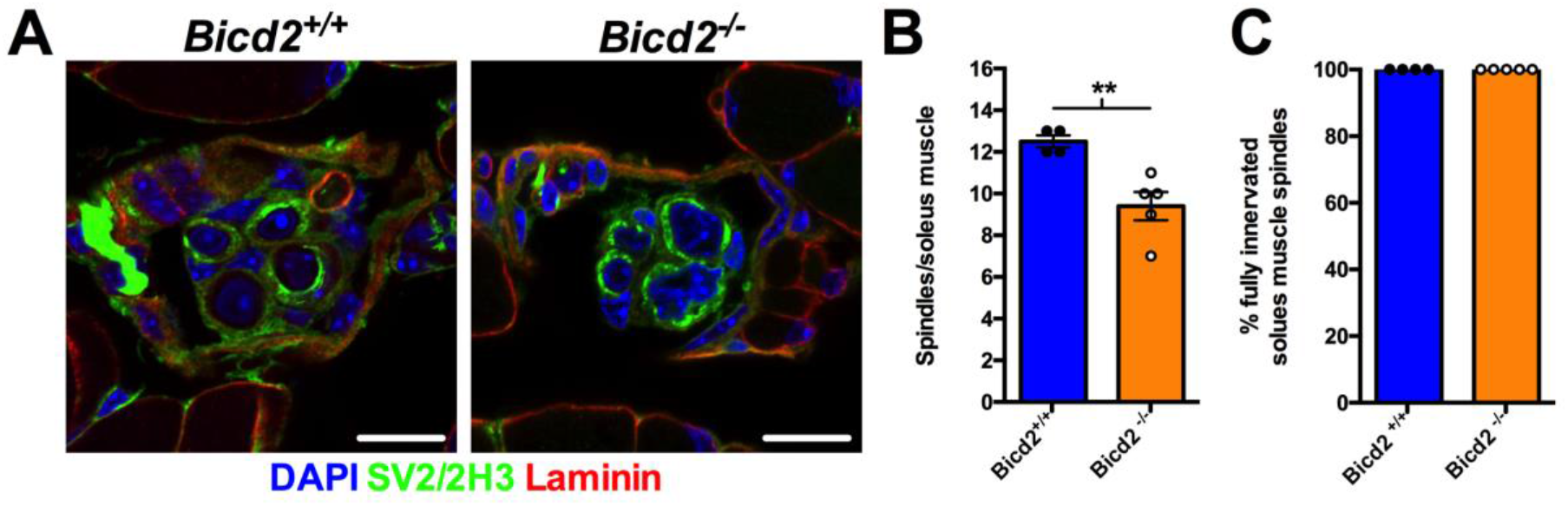
Loss of muscle spindles in *BICD2*^−/−^ mice at 21 days of age. (**A**) shows example images of a cross section through a muscle spindle in *Bicd2*^+/+^ and *BICD2*^−/−^ mice stained for nuclei (DAPI, blue), the neuronal marker SV2/2H3 (green) and laminin (muscle membrane, red). Scale bars = 10 μm. (**B**) shows the total number of muscle spindles in the soleus muscle of *Bicd2*^+/+^ and *BICD2*^−/−^ mice, \*\**p*=0.0065 (unpaired *t*-test, n=4-5). (**C**) shows full innervation patterns in the muscle spindles of *Bicd2*^+/+^ and *BICD2*^−/−^ mice. Error bars = SEM.

### Patient fibroblasts show evidence of impaired secretion

Rab6, a small GTPase interacting with BICD2, is an established regulator of the secretory pathway (Grigoriev et al., 2007; Fourriere et al., 2019) and controls the flow of secreted proteins that are transported from the Golgi to the plasma membrane via a microtubule-dependent process. As a consequence, loss of Rab6 results in a global reduction of protein secretion (Bardin et al., 2015).

We therefore hypothesised that SMALED2-causing mutations in *BICD2* result in an impairment of Rab6 function through its mislocalisation, impairing the targeting of secretory vesicles to the plasma membrane. This would explain why mutations in the N-terminal domain of BICD2, which enhance retrograde processive motility leading to accumulation of BICD2 at the centromere (Huynh and Vale, 2017; Peeters et al., 2013), and C-terminal mutations that impair the recruitment of Rab6, but not dynein binding, to BICD2 cause an identical phenotype (Rossor et al., 2015).

To verify this hypothesis, we performed an established VSV-G secretion assay (Miserey-Lenkei et al., 2010) using fibroblasts from a patient with a *BICD2*^I189F^ SMALED2 mutation and from an age-matched control (Rossor et al., 2015). These cells were transfected with a plasmid encoding for a temperature-sensitive mutant of vesicular stomatitis virus glycoprotein (ts0-45 VSV-G) tagged with GFP and incubated at 40°C for 14 h. This high temperature treatment results in misfolding of ts0-45 VSV-G and its retention in the ER. Cooling of the cells to 32°C allows the refolding of VSV-G and its targeting to the plasma membrane, which could then be quantified. This assay revealed a significant reduction in the rate of secretion of VSVG over time (**Figure 5**). Similarly to previous experiments performed in Rab6 knockout cell lines (Grigoriev et al., 2007; Miserey-Lenkei et al., 2010), secretion was significantly delayed, but not abolished. To confirm that this difference was not simply due to clonal differences, the accumulation of VSV-G at the plasma membrane at 240 minutes (the time point with the largest difference) was repeated for two additional, unrelated, age-matched controls and SMALED2 patients bearing the S107L and R501P mutations in BICD2. These additional experiments showed a significant reduction in VSV-G appearance on the plasma membrane at 240 minutes (**Figure 5**; control mean = 0.64±0.017, n=3; SMALED2 = 0.52±0.043, n=3; unpaired *t*-test *p*=0.05).

**Figure 5.**
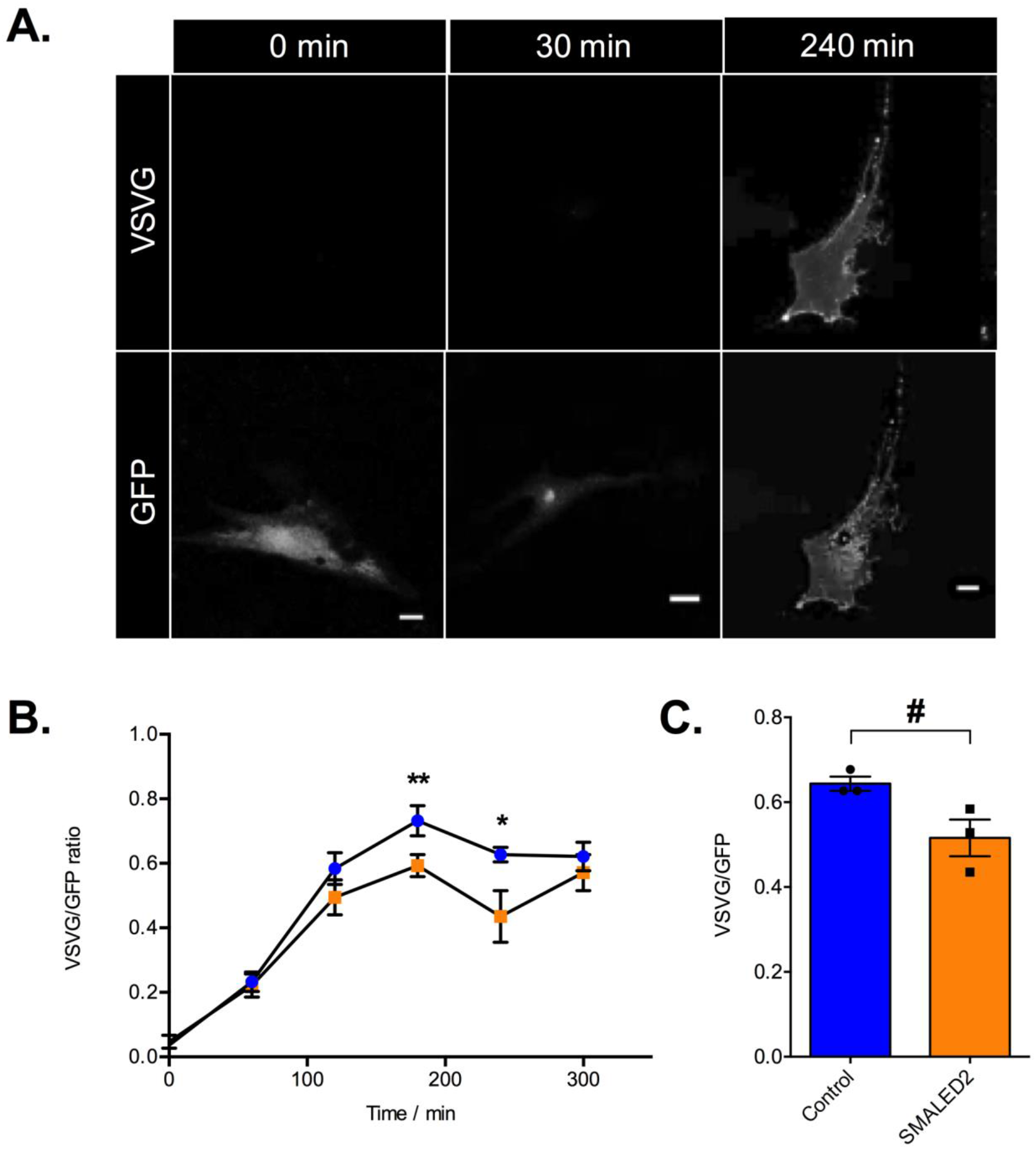
SMALED2 patient fibroblasts show delayed VSV-G secretion compared to controls. (**A**) is an example of a human fibroblast transfected with a plasmid encoding for the thermo-sensitive vesicular stomatitis virus glycoprotein (ts0-45 VSVG) at 32ºC. The time prior to fixation is indicated on the top; staining with an anti-VSV-G [8G5F11] against a surface epitope of VSV-G in non-permeabilised cells (top row of panels); GFP staining is shown in the bottom row. Scale bars = 20 μm. (**B**) Kinetics of VSV-G secretion in fibroblasts isolated from a patient with SMALED2 (I189F mutation, orange) and an age-matched control (blue) are quantified as the ratio of total surface VSV-G staining to total GFP. The x axis shows the time in minutes at 32ºC prior to fixation (n=10 cells per condition; \*\**p*=0.008, \**p*=0.009; multiple unpaired *t*-tests corrected for multiple comparisons using the Holm-Sidak method). (**C**) shows the average ratio of surface VSV-G to total GFP at 240 min for three independent healthy control and three unrelated SMALED2 (S107L, I189F and R501P) fibroblast cell lines (# p=0.052, unpaired t-test). Error bars = SEM.

Since BICD2 and Rab6 have also been involved in COPI-independent transport from the Golgi to the ER (White et al., 1999; Matanis et al., 2002), we investigated the retrograde flow from the Golgi to the ER using galactosyltransferase–GFP in Brefeldin A-treated control and SMALED2 fibroblasts, but did not detect any significant difference between patient and control cells (data not shown).

In light of these results, we propose a non-cell autonomous mechanism of motor neuron loss in SMALED2, which is also likely to be applicable to SMALED caused by mutations in *DYNC1H1*. We have shown that loss of BICD2 leads to death of motor neurons and that this is a non-cell autonomous process driven by the loss of BICD2 in muscle. Altogether, our results demonstrate that the *BICD2*^−/−^ mouse is a good model of SMALED2 as *Bicd2*^−/−^ mice and patients with severe SMALED2 mutations show similar clinical features including hydrocephalus, cerebellar hypoplasia and cortical migration defects (Ravenscroft et al., 2016; Jaarsma et al., 2014a). A non-cell autonomous pathomechanism is also supported by a previous study demonstrating that the cerebellar hypoplasia in *Bicd2*^−/−^ is driven by non-cell autonomous deficits arising in Bergmann glia (Jaarsma et al., 2014).

Quantitative analyses of secretion in patient fibroblasts compared to controls shows only a modest secretion deficit, as found in Rab6 knockout models (Bardin et al., 2015). How such a delay in secretion causes a significant loss of motor function can be explained in light of the neurotrophin hypothesis in which a surplus of motor neurons compete for muscle-secreted neurotrophic factors for survival (Kanning et al., 2010). In this model, a reduction in the rate of secretion would yield a decrease in neurotrophin availability, driving an excess of motor neuron death. This hypothesis predicts that no progressive motor neuron degeneration should occur in SMALED2 and muscle-deficient *BICD2* mice, as demonstrated in this work and elsewhere (Oates et al., 2012; Rossor et al., 2015).

Our model may also explain why the *loa*, *crawling* and *sprawling* mouse models, which bear point mutations in *Dync1h1* develop gamma motor neuron and sensory 1a afferent loss (Ilieva et al., 2008; Chen et al., 2007; Hafezparast et al., 2003). Both gamma motor neurons and 1a sensory afferents innervate muscle spindles and rely on muscle-derived neurotrophic factor secretion during early development. Therefore, an impairment of localised neurotrophin secretion from the muscle of *dync1h1* mutant muscles would lead to a loss of gamma motor and sensory 1a neurons.

Our model does not align with previous results generated using a *Drosophila* model of SMALED2 (Martinez Carrera et al., 2018), which led to the proposal that this pathology is determined by a cell autonomous effect of mutant *BICD2* in motor neurons (Martinez Carrera et al., 2018). However, *Drosophila* have only a bicaudal D protein (BicD), unlike mice and humans, which express four bicaudal D proteins (BICD1, BICD2, BICDR1 and BICDR2) with different functions and interacting partners (Hoogenraad and Akhmanova, 2016; Terenzio and Schiavo, 2010). As such, the model of Martinez-Carrera is expected to be unable to differentiate between the effects of BICD1 and 2, which is crucial, as we have previously shown that BICD1 has an essential cell autonomous role in the sorting of signalling endosomes in motor neurons (Terenzio et al., 2014a; b). Furthermore, *Drosophila* display important differences to mammals in the neurotrophin pathway at the NMJ (Sutcliffe et al., 2013).

Our model provides strong evidence in support of a non-cell autonomous mechanism of motor neuron loss in SMALED2 which involves impaired secretion of muscle-derived neurotrophins during development. However, a number of important questions still need to be addressed in future studies. The nature of the secreted muscle-derived neurotrophic factor(s) is still unclear as well as whether the phenotypes of mice with loss-of-function mutations in *Bicd2* are similar to that of *Bicd2*^−/−^ mice. However, the most important point to clarify in future studies is likely to be the precise time point at which motor neuron loss occurs, which is important when considering future therapeutic interventions in SMALED2 patients.

## Materials and Methods

### Animals

Animal experimentation was performed under license from the United Kingdom Home Office in accordance with the Animals (Scientific Procedures) Act (1986), and approved by the University College London Queen Square Institute of Neurology ethics committee. Mice homozygous for an allele in which the loxP sequence is inserted into intron 1 and the 3’UTR of the endogenous *Bicd2* gene (Jaarsma et al., 2014) were crossed with heterozygous deleter-Cre mice (C57BL/6NTac-*Gt(ROSA)26Sor^tm16(cre)Arte^*)(Otto et al., 2009) to generate *Bicd2* heterozygous knockout mice expressing a Cre-recombinase transgene. These mice were subsequently crossed with C57Bl/6J mice to generate *Bicd2*^+/−^ mice with no Cre recombinase transgene. Genotypes of mice were determined by PCR of ear clip DNA (P1 = AATGGAGAAGATCTCATCTTGGCAGG, P2 = GTGTAGCACTTCAGGAACATCCATGC, P3 = TGTCAGCAAACTCCATCTCTAGCCTC, P261 = CGGCGGCATCAGAGCAGCCGATTG).

To generate muscle-specific *Bicd2* knockout mice, homozygous *Bicd2*^*loxP*/*loxP*^ mice (C57Bl/6J) were crossed with knockin *MyoD-Cre* mice (FVB.Cg-*Myod1^tm2.1(icre)Glh^*/J, Jackson laboratory) (Kanisicak et al., 2009) and back-crossed to the original *Bicd2*^*loxP*/*loxP*^ progeny for six generations to produce a congenic strain. To generate motor neuron-specific *Bicd2* knockout mice, homozygous *Bicd2*^*loxP*/*loxP*^ mice (C57Bl/6J) were crossed with *ChAT-IRES-Cre* knock-in mice (B6;129S6-*Chat^tm2(cre)Lowl^*/J, Jackson laboratory) (Rossi et al., 2011) and then back-crossed to the original *Bicd2*^*loxP*/*loxP*^ progeny to generate a homozygous *Bicd2*^loxP/loxP^ background.

### Histopathology

Mice were terminally anaesthetised with intraperitoneal injection of pentobarbitone followed by thoracotomy and transcardial perfusion with 10 ml 0.9% NaCl followed by 20 ml 4% paraformaldehyde (PFA, Fisher Scientific) in phosphate buffered saline (PBS). L4 DRG and L4 dorsal and ventral roots were dissected from PFA-fixed mice and post-fixed in PFA/glutaraldehyde buffer at 4°C (20 ml 0.9% saline, 10 ml 10% PFA, 10 ml 10% glutaraldehyde (Sigma-Aldrich), 20% dextran (20,000 MW, Sigma-Aldrich) made up to 100 ml with 0.1 M PIPES-NaOH, pH 7.6, for 24 h before fixation in 1% osmium tetroxide (Agar Scientific) and processing into araldite CY212 epoxy resin (Agar Scientific) through graded alcohols and propylene oxide solutions using a standard protocol. Semi-thin sections (0.8 μm) were cut on an Ultracut E ultramicrotome (Leica), stained with 1% toluidine blue containing 1% borax (BDH), and examined with a Leica light microscope using an oil immersion 100x lens. Images were generated by image stitching using HUGIN-panorama photo stitching. Axon diameter was calculated using an in-house semi-automated thresholding programme using Definiens image analysis software.

### Immunohistochemistry

#### Muscle fibre subtype and area quantification

P21 mice were euthanized by cervical dislocation and skeletal muscles (gastrocnemius) were dissected and snap frozen in liquid nitrogen-cooled isopentane and stored at −80°C for cryosectioning. Immunofluorescent staining was carried out on 10 μm frozen sections. Sections were blocked in PBS 0.2% Triton X-100, 5% goat serum for 1 h. Primary and secondary antibodies were diluted in PBS 0.2% Triton X-100, 2% goat serum. Primary antibodies: Type 1 BA-D5 (mouse IgG2b) DSHB (1 in 100), Type IIA SC-71 (mouse IgG1) DSHB (1 in 100), Type IIB BF-F3 (mouse IgM) DSHB (1 in 100) and laminin (rabbit) L9393 Sigma-Aldrich (1 in 100) were incubated for 1 h at 37°C. Sections were then washed three times with PBS followed by goat anti-mouse IgG2b conjugated to AlexaFluor488, goat anti-mouse IgG1 conjugated to AlexaFluor647, goat anti-mouse IgM conjugated to AlexaFluor568 and goat anti-rabbit conjugated to AlexaFluor410 (all from Life Technology, 1:500). Secondary antibodies were incubated for 30 min at 37°C and washed three times with PBS. Cover slips were mounted on stained sections using Dako fluorescent mounting media and dried for 48 h. Z-stack stitched images were acquired using a Zeiss LSM 710 confocal microscope using previously described parameters (Mayeuf-Louchart et al., 2018). Quantification of the fibre number, area and type were performed using the muscle-J automated freeware and Image J as previously described (Mayeuf-Louchart et al., 2018).

#### NMJ analyses

Muscles were processed for immunohistochemistry and NMJ phenotypes scored as previously described (Sleigh et al., 2014). 100 NMJs were scored per mouse for poly-innervation and occupancy counts, whilst 16-20 NMJs were used for area measurements.

#### Muscle spindle assessment

Soleus muscles were dissected and processed, and spindles analysed as described previously with minor modifications (Sleigh et al., 2017). Briefly, soleus muscles were dissected then fixed overnight in 4% PFA in PBS prior to overnight equilibration in 20% (w/v) sucrose in PBS before freezing in Tissue-Tek O.C.T. (Sakura Finetek). 20 μm transverse serial sections throughout the entire muscle were cut onto three parallel slides for immunohistochemistry and analyses using an OTF Cryostat (Bright Instruments)

#### DRG dissection and staining

Mice were perfused with 4% PFA in PBS before L4 DRG were dissected as previously described (Sleigh et al., 2016). DRG were sectioned and stained as previously published (Sleigh et al., 2017). Briefly, DRG were post-fixed overnight in 4% PFA in PBS, before embedding in Tissue-Tek O.C.T., and sectioning at 10 μm across four parallel polysine-coated slides (VWR, 631-0107) with an OTF Cryostat. DRG sections were permeabilised for 30 min in PBS containing 0.3% Triton X-100 and blocked for 30 min in 10% bovine serum albumin (BSA) and 0.3% Triton X-100 in PBS, before probing overnight at 4°C with primary antibodies (1:500 mouse anti-NF200 [N0142; Sigma-Aldrich] and 1:500 rabbit anti-peripherin (AB1530; Merck Millipore) in blocking solution. Sections were then washed with PBS for 30 min, probed for 2 h with secondary antibodies (1:1,000) in PBS, before washing in PBS, flooding with fluorescence mounting medium (S3023; Dako) and covering with a 22 × 50 mm cover glass (VWR).

### VSVG assay

Frozen vials of human skin fibroblasts were thawed and grown in T75 flasks for seven days in DMEM media supplemented with 20% foetal calf serum and L-glutamine (all from Gibco). Cells were trypsinised and plated onto 13 mm untreated glass coverslips (50,000 per coverslip). After 24 h, fibroblasts were transfected with a GFP-VSV-G ts045 plasmid using Lipofectamine 3000 (Invitrogen) as per manufacturer’s instructions and grown at 37°C. After 8 h, cells were incubated at 40°C for 14 h before being transferred to an incubator at 32°C. Cells were fixed at 0, 60, 120, 180, 240 and 300 min after incubation at 32°C in PFA 4% in PBS. Non-permeabilised cells were then incubated with PBS 1% BSA for 15 min prior to overnight incubation with mouse anti-VSV-G [8G5F11; Kerafast] antibody (1:2000) at 4°C. Cells were washed three times with PBS before incubation with a donkey anti-mouse AlexaFluor568 secondary antibody (Life Technology, 1:500) for 1 h at room temperature. Cells were then washed, stained with DAPI (1:2,000) and coverslips mounted and fixed with DAKO fluorescent mounting media. Ten fibroblasts per condition were imaged with a Zeiss LSM 510 confocal microscope and the ratio between plasma membrane VSV-G to the total VSV-G (GFP signal) was quantified.

### Brefeldin A assay

Human fibroblasts were cultured as previously described above, but were plated on 20 mm coverslips in 6 well plates at a density of 100,000 per well. After 24 h in culture, cells were transfected with a plasmid encoding human galactosyltransferase tagged with GFP (GT–GFP) (kind gift from Masayuki Murata, Tokyo). 24 h post-transfection, cells were imaged using a Zeiss LSM 710 confocal microscope. Briefly, cells were imaged at 37°C, 5% CO_2_ with a 40x objective. The Golgi apparatus was identified by GFP staining. Brefeldin A (10 μg/ml final concentration; Sigma-Aldrich) was re-suspended in DMEM, 10% FCS, 20 mM HEPES-NaOH (final concentration), pH 7.3, and added to the coverslip at time 0. Time-lapse imaging at 5 s intervals was carried out and the time recorded to the beginning and end of the ‘Golgi blush’, defined as the stereotypical morphological changes in Golgi morphology leading to its resorption into the ER.

### Statistical analysis

Data were assumed to be normally distributed unless evidence to the contrary was provided by the D’Agostino and Pearson omnibus normality test. Data were statistically analysed using an unpaired *t* test, Mann-Whitney *U*-test, or one-way ANOVA with post-hoc Dunnett’s test against a control mean. GraphPad Prism 6 software was used for all statistical analyses. Means ± standard error of the mean (SEM) were plotted for all graphs.

## Online supplemental material

**Figure S1** shows a comparison of the ratio of NF200 and peripherin positive sensory neurons in the L4 DRG of *Bicd2*^+/+^ and *Bicd2*^−/−^ mice. **Figure S2** shows a quantitative comparison of muscles and muscle fibre type in *Bicd2*^+/+^ and *Bicd2*^−/−^ mice. **Figure S3** shows the haematoxylin and eosin staining, transmission electron microscopy and immunohistochemical staining of muscle fibre types in the gastrocnemius muscle of p21 *Bicd2*^+/+^ and *Bicd2*^−/−^ mice.

## Acknowledgements

We are grateful to Professor Kathryn North and Dr Emily Oates for providing patient fibroblasts, to the MRC Centre Biobank in London for provision of the patient fibroblast lines used in this study and to Pierpaolo Ala, technician in the Biobank. The Biobank is also supported by the National Institute for Health Research (NIHR) Great Ormond Street (GOS) Hospital Biomedical Research Centre. We are also grateful to Stéphanie Miserey-Lenkei for providing the GFP-VSV-G ts045 plasmid and for technical advice. We are also grateful to Professor Masayuki Murata for providing the GT-GFP plasmid. We are also grateful to Professor Rob Brownstone and Dr Nadine Simons-Weidenmaier for providing the ChAT-Cre mice and to Dr Stephane Nedelec for critical reading of the mnuscript. J.N.S. is funded by the Medical Research Council Career Development Award (MR/S006990/1). This work was supported by a Wellcome Trust Senior Investigator Award 107116/Z/15/Z (to G.S.), the European Union’s Horizon 2020 Research and Innovation programme under grant agreement 739572 (to G.S.), a UK Dementia Research Institute Foundation award (to G.S.). and a Wellcome Trust Postdoctoral Fellowship for Clinicians (110043/Z/15/Z) (A.M.R). The core support of MDUK to the UCL Neuromuscular Centre is also gratefully acknowledged

## Author contributions

A.M.R. wrote the draft of the manuscript and performed dissections, cell culture experiments and data analysis. A.M.R. and G.S. designed the experiments. J.N.S. performed the NMJ, muscle spindle and DRG analyses. MG produced the semi-thin nerve root sections. C.H. provided the *Bicd2*^*loxP/loxP*^mice and F.M. and M.M.R. provided essential reagents. A.M.R., J.N.S. and G.S. revised the text and figures. The authors declare no competing financial interests.

## Abbreviations

SMALED: Spinal Muscular Atrophy Lower Extremity Predominant
BICD2: Bicaudal D2
VSV: Vesicular Stomatitis Virus
BFA: Brefeldin A

