## Supplementary figure for "Loss of BICD2 in muscle drives motor neuron loss in a developmental form of spinal muscular atrophy"

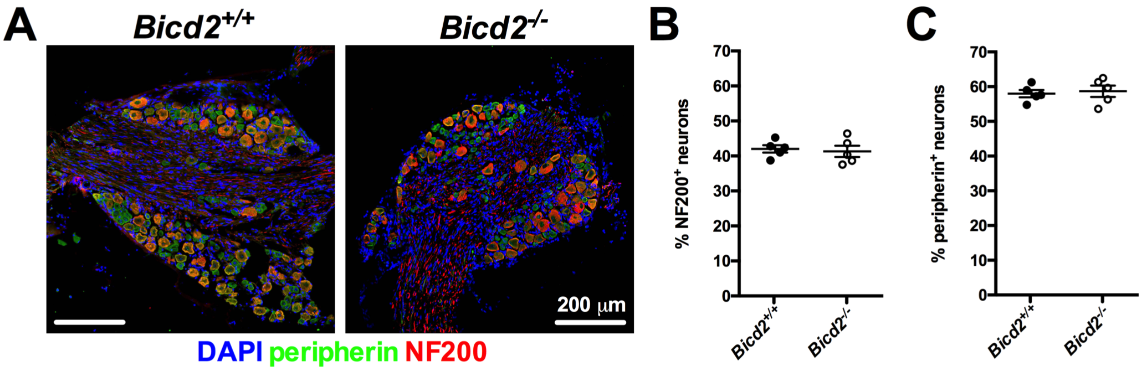


**Supplementary Figure S1**. There is no difference in the percentages of medium-to-large (NF200$^+$) and small (peripherin$^+$) sensory neurons between *Bicd2*$^{+/+}$ and *Bicd2*$^{-/-}$ mice. (**A**) shows representative images of L4 dorsal root ganglia stained for DAPI (blue), peripherin (green) and NF200 (red). (**B & C**) There is no difference in the percentages of NF200$^+$ or peripherin$^+$ sensory neurons between *Bicd2*$^{+/+}$ and *Bicd2*$^{-/-}$ mice (NF200, *Bicd2*$^{+/+}$ 42% ±1 (n=5), *Bicd2*$^{-/-}$ 41% ±1.6, unpaired *t*-test p=0.73, peripherin, *Bicd2*$^{+/+}$ 58% ±1 (n=5), *Bicd2*$^{-/-}$ 59% ±1.6, unpaired *t*-test p=0.73).

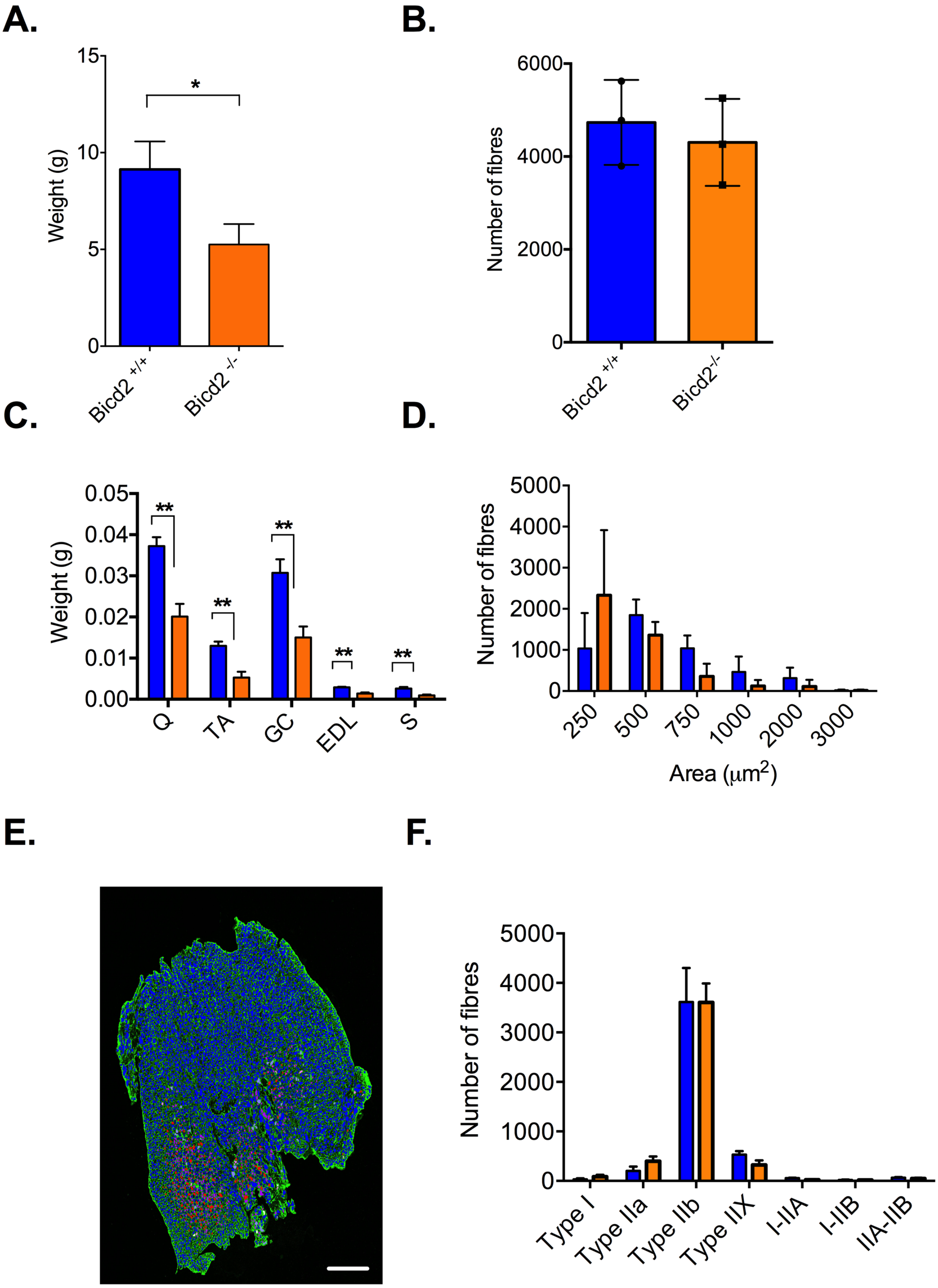
A.
B.
C.
D.
E.
F.
Weight (g)
Bicd2 +/+
Bicd2 -/-
Number of fibres
Bicd2 +/+
Bicd2-/-
Q
TA
GC
EDL
S
Area (μm2)
Type I
Type IIa
Type IIb
Type IIX
I-IIA
I-IIB
IIA-IIB

**Supplementary Figure S2**. A quantitative comparison of the gastrocnemius muscle in *Bicd2*$^{+/+}$ and *Bicd2*$^{-/-}$ mice at 21 days of age. (**A**) shows the weight in grams of *Bicd2*$^{+/+}$ (n=5) and *Bicd2*$^{-/-}$ (n=4) mice, *$p$=0.0028 (unpaired *t*-test). (**B**) shows the number of muscle fibres of the gastrocnemius muscle in *Bicd2*$^{+/+}$ (n=3) and *Bicd2*$^{-/-}$ (n=3) mice. (**C**) shows a comparison of the weight in grams in the muscles of *Bicd2*$^{+/+}$ (blue, n=3) and *Bicd2*$^{-/-}$ (orange, n=3) mice. Q=quadriceps, TA=tibialis anterior, GC=gastrocnemius, EDL=extensor digitorum longus, S=soleus, **$p$<0.01 (multiple t-tests corrected for multiple comparisons using the Holm-Sidak method). (**D**) is a histogram of the gastrocnemius muscle fibres according to diameter (blue=*Bicd2*$^{+/+}$, orange=*Bicd2*$^{-/-}$). (**E**) shows a digital reconstruction generated from images presented in **Supplementary Figure 3F** following automated fibre type quantification using an ImageJ plugin called 'muscle-J'. Red = type 1, magenta = type IIA, dark blue = type IIX, light blue = type IIB). (**F**) shows the mean number of muscle fibre types in the gastrocnemius muscle of *Bicd2*$^{+/+}$ (n=3) and *Bicd2*$^{-/-}$ (n=3) mice. Error bars = SEM.

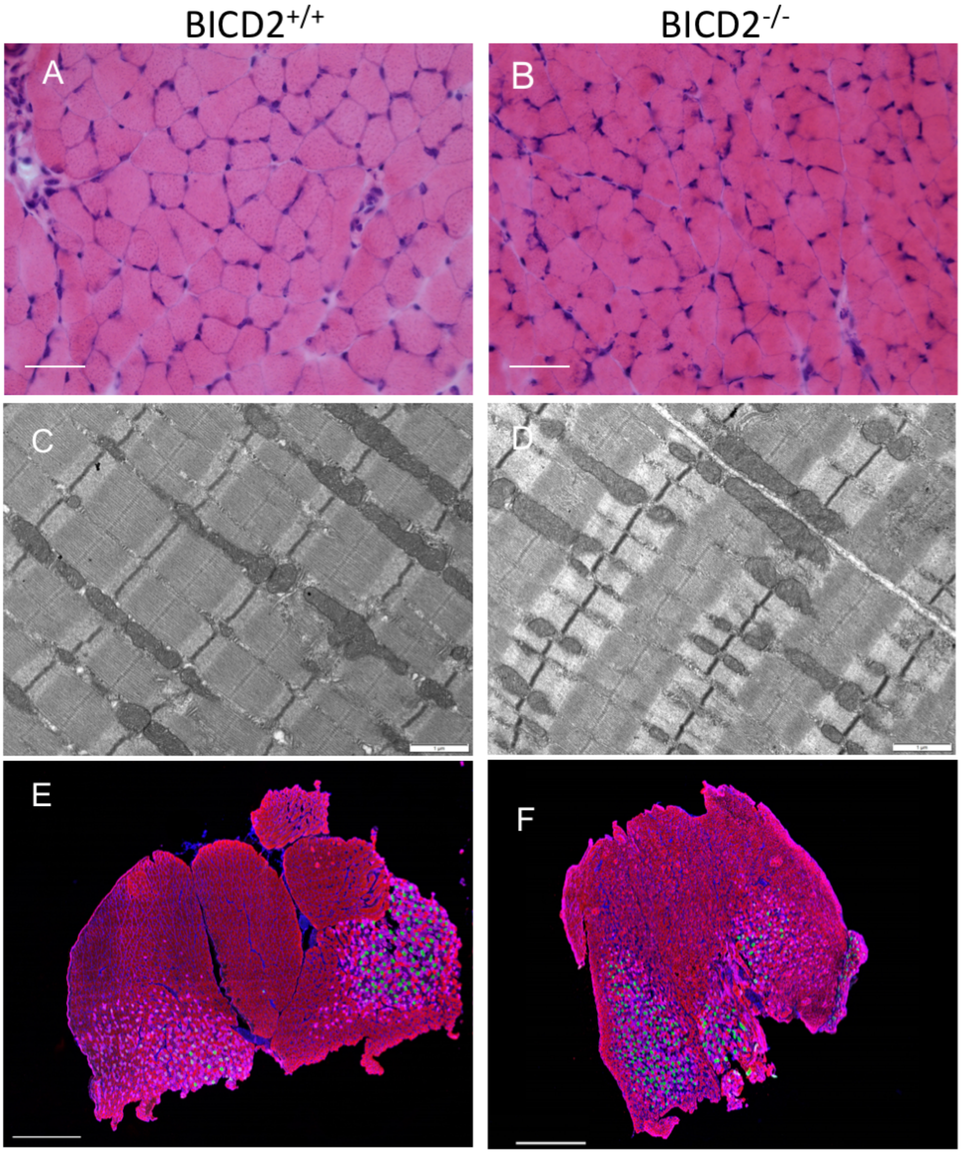


**Supplementary Figure S3**. No histological differences between *Bicd2*$^{+/+}$ and *Bicd2*$^{-/-}$ gastrocnemius muscle at 21 days of age. Haematoxylin and eosin stains of the gastrocnemius muscle of *Bicd2*$^{+/+}$ (**A**) and *Bicd2*$^{-/-}$ (**B**) mice, scale bars = 50 µm. Transmission electron microscopy image of the gastrocnemius muscle in *Bicd2*$^{+/+}$ (**C**) and *Bicd2*$^{-/-}$ (**D**) mice. Scale bars = 1 µm. (**E** & **F**) show immunohistochemical staining for muscle fibre types in 10 µm cryosections of the gastrocnemius muscle in *Bicd2*$^{+/+}$ and *Bicd2*$^{-/-}$ mice, respectively. Green= type 1, pink = type IIa, red = type IIb, no staining = type IIx, blue = laminin, scale bars = 500 µm.
